# MIR-128 TARGETING PPAR-γ IMPROVES VASCULAR REMODELING IN SPONTANEOUSLY HYPERTENSIVE RATS BY REGULATING THE TLR4/NFKB INFLAMMATORY PATHWAY

**DOI:** 10.64898/2026.05.05.723109

**Authors:** Zhoufei Fang, Han Cai, Ruonan Liu, Lele Yu, Chunxian Chen, Shikun Chen, Lirong Li, Qiujing Chen, Huange Cai, Jinzi Su, Feng Peng

**Affiliations:** Department of Geriatrics, the First Affiliated Hospital of Fujian Medical University, Fujian Medical University, FujianFuzhou, 350004, People’s Republic of China; Department of Cardiology, Binhai Campus of the First Affiliated Hospital, National Regional Medical Center, Fujian Medical University, Fuzhou, 350212, People’s Republic of China; Department of Cardiology, the First Affiliated Hospital of Fujian Medical University, Fuzhou, Fujian 350004, People’s Republic of China; The First Affiliated Hospital, Fujian Hypertension Research Institute, Fujian Medical University, Fuzhou, Fujian, 350004,People’s Republic of China; Clinical Research Center for Geriatric Hypertension Disease of Fujian province, The First Affiliated Hospital of Fujian Medical University, Fuzhou, People’s Republic of China; Clinical Research Center for Metabolic Heart Disease of Fujian Province, the First Affiliated Hospital, Fujian Medical University, Fuzhou, Fujian, China; The Higher Educational Key Laboratory for Cardiovascular Disease of Fujian Province, the First Affiliated Hospital, Fujian Medical University, Fuzhou, Fujian, China; Department of Geriatrics, National Regional Medical Center, Binhai Campus of the First Affiliated Hospital, Fujian Medical University, Fuzhou, Fujian, China; Clinical Research Center for Geriatric Hypertension Disease of Fujian province, The First Affiliated Hospital of Fujian Medical University, Fuzhou, Fujian, China; Branch of National Clinical Research Center for Aging and Medicine, The First Affiliated Hospital of Fujian Medical University, Fuzhou, Fujian, China; Department of Ultrasound, the First Affiliated Hospital of Fujian Medical University, Fuzhou, Fujian 350004,People’s Republic of China

**Keywords:** hypertension, renal denervation (RDN), PPAR-γ/TLR4/NF-κB axis, microRNA-128 (miR-128)

## Abstract

**OBJECTIVE:** This study aimed to explore the role and underlying mechanism of microRNA-128 (miR-128) in regulating vascular remodeling in spontaneously hypertensive rats (SHRs), focusing on its targeting of peroxisome proliferator-activated receptor γ (PPAR-γ) and modulation of the Toll-like receptor 4/nuclear factor-κB (TLR4/NF-κB) inflammatory pathway.

**METHODS:** All experimental procedures were approved by the Animal Care and Use Committee of Fujian Medical University. In vivo, ten-week-old male SHRs were randomly assigned to three groups: renal denervation (RDN, n=6), sacubitril/valsartan (Sac/Val, n=6), and Sham (n=6). Age-matched Wistar-Kyoto (WKY) rats served as normotensive controls (n=6).Eight weeks after intervention, mesenteric arteries were harvested for histological, functional, and molecular analyses. Serum miR-128 levels were measured by quantitative real-time polymerase chain reaction (qRT-PCR). The expression levels of key proteins in the vascular wall were assessed via immunofluorescence (IF), immunohistochemistry (IHC), and Western blotting (WB). Bioinformatics analysis and RNA sequencing (RNA-seq) were employed to identify core genes and signaling pathways associated with hypertension-induced pathological inflammation.

**RESULTS:** In vivo, in the SHR sham-operated group, elevated blood pressure, severe vascular remodeling, and impaired vasodilatory function were observed, accompanied by downregulated miR-128 expression and upregulated TLR4/NF-κB signaling activity (all p < 0.0001).RDN postoperative, miR-128 expression was significantly restored, which in turn inhibited the TLR4/NF-κB pathway, reduced the production of pro-inflammatory cytokines (including IL-1β, IL-6, and TNF-α), and ameliorated vascular dilation dysfunction in SHRs (all p < 0.0001). Mechanistically, miR-128 negatively regulated the TLR4/NF-κB signaling pathway while upregulating the expression of PPAR-γ (p < 0.05).

**CONCLUSION:** RDN not only exerts a hypotensive effect but also improves hypertensive vascular remodeling. miR-128 inhibits excessive inflammation in vascular smooth muscle cells and alleviates vascular remodeling in SHRs via the PPAR-γ/TLR4/NF-κB axis. These findings identify miR-128 as a potential therapeutic target for RDN in the treatment of hypertension, providing a novel regulatory strategy for the precision management of cardiovascular diseases.

## 1 **Introduction**

Hypertension represents the leading global risk factor for cardiovascular morbidity and mortality[1]. Vascular remodeling, characterized by excessive vascular smooth muscle cell proliferation, migration, extracellular matrix deposition, and persistent vascular inflammation, is a core pathological process driving the progression of hypertension and subsequent target organ damage[2]. Despite advances in pharmacological management, poor adherence and incomplete blood pressure control remain major clinical challenges[3].

Renal denervation (RDN) is a minimally invasive intervention that reduces systemic sympathetic overactivity by ablating renal afferent and efferent sympathetic nerves[4]. Accumulating evidence demonstrates that RDN exerts beneficial effects beyond blood pressure reduction, including attenuation of vascular remodeling and suppression of vascular inflammation[5,6]. However, the precise molecular mechanisms mediating these vascular protective effects remain incompletely defined.

MicroRNAs (miRNAs) are highly conserved small non-coding RNAs that regulate gene expression post-transcriptionally[7]. MiR-128 is abundantly expressed in cardiovascular tissues and participates in the regulation of cell proliferation, apoptosis, and inflammatory responses[8,9]. Our preliminary studies revealed dysregulated miR-128 expression in hypertensive animal models, with a strong correlation to vascular inflammation and remodeling severity. Bioinformatic predictions further suggested a potential targeting relationship between miR-128 and PPAR-γ[10].

PPAR-γ is a nuclear receptor with well-established anti-inflammatory, anti-fibrotic, and vascular-protective properties[11]. It negatively regulates the TLR4/NF-κB signaling cascade, a central mediator of vascular inflammation in hypertension[12]. Based on these observations, we hypothesized that RDN improves vascular remodeling in hypertension by modulating the miR-128/PPAR-γ/TLR4/NF-κB axis[13,14].

This study investigated the effects of RDN on miR-128 expression, vascular structure, endothelial function, and inflammatory signaling in SHRs. We demonstrate that miR-128 directly targets PPAR-γ to regulate TLR4/NF-κB-mediated inflammation, thereby contributing to the therapeutic benefits of RDN. These findings provide novel mechanistic insights into RDN and identify a potential molecular target for the treatment of hypertensive vascular remodeling.

## 2 Material and Methods

All experimental protocols were approved by the Animal Care and Use Committee of Fujian Medical University. Ten-week-old male ( at baseline) SHRs and age-matched WKY rats were purchased from Vital River Laboratory Animal Technology Co., Ltd. (Beijing, China). SHRs were randomly divided into three groups: Sham surgery (Sham), renal denervation (RDN), and sacubitril/valsartan (Sac/Val) treatment. WKY rats served as normotensive controls.All assays were conducted when rats were

18 weeks of age (Intervention endpoint=8 weeks after intervention).

### 2.1 Surgical Procedures

Following a one-week period of adaptive feeding, 10-week-old SHR were administered RDN under intraperitoneal anesthesia with urethane (800 mg/kg) and

α-chloralose (40 mg/kg). A surgical incision was made in the lateral posterior flank of the peritoneum, allowing for the kidney to be exposed. The surrounding organs of the renal artery and vein were protected with saline infiltrated cotton, while a small piece of cotton soaked in capsaicin solution (33 mM dissolved in 5% ethanol, 5% tween and 90% saline) was tightly wrapped around the renal artery and vein for ten minutes.

Following the treatment duration, the cotton was removed, saline was used to cleanse the affected area, and the wound was sutured. The same procedure was applied to the contralateral side.

### 2.2 Measurement of Blood Pressure

The conscious rats underwent systolic blood pressure (SBP) and heart rate (HR) monitoring utilizing a noninvasive tail-cuff system (BP-300, Chengdu Taimeng Science and Technology Company, China). To register the pulsations of the tail artery, the rats were kept in a thermostatic chamber at 30℃ for 10 minutes before each measurement. Averaged from three readings, the values were recorded at baseline (10 weeks of age) and every 2 weeks thereafter until the conclusion of the experiment (18 weeks of age).

### 2.3 Assessment of Arterial Baroreflex Function

In this study, rats were subjected to intraperitoneal injection of ethyl carbamate (800 mg/kg) and α-chloralose (40 mg/kg) for anesthesia prior to the insertion of polyethylene catheters (PE-50) into the carotid artery and femoral vein. The carotid artery was then cannulated and connected to a pressure transducer for blood pressure recording, while the femoral vein was connected for drug administration. The upper limb was affixed with a II-lead ECG to record heart rate. Blood pressure and heart rate were measured independently using the BL-420S Biofunctional Experiment System (Chengdu Taimeng Science and Technology Company, China). At the conclusion of the cannulation process, arterial baroreflex sensitivity (BRS) measurements were taken 30 minutes after to evaluate arterial baroreflex function. In addition, phenylephrine (PE: 8μg/kg) was administered via the femoral vein to provoke an acute increase in blood pressure and reflexively decelerate HR, and changes in mean arterial pressure (ΔMAP) andΔHR were recorded pre-and post-drug administration. (BRS =ΔHR/ΔMAP).

### 2.4 Vascular Histopathological Examination

Expose the abdominal aorta for blood collection and intubation; The blood vessels were dilated with 0.1mg/mL sodium nitroprusside and fixed by perfusion with a 10% formaldehyde solution containing eosin under a column pressure of 60-80cmH2O for 12 hours. Mesenteric arteries were taken, fixed with 4% paraformaldehyde, subjected to gradient dehydration, paraffin-embedded, and continuously sectioned.

Hematoxylin-eosin (HE) staining was performed respectively With Masson staining. The vascular wall thickness/lumen radius (WT/LR), vascular wall area/lumen area (W/L), and collagen fiber content were analyzed by Image J software to reflect the degree of vascular remodeling.

### 2.5 ELISA for Norepinephrine (NE) Levels

100mg samples were taken from renal cortex and added to 10ml PBS solution. The mixture was electrically homogenized under an ice bath at a rate of 7000 r/min for 15 seconds, and this procedure was repeated thrice. The obtained homogenate was then centrifuged at a rate of 3000 r/min for 20 minutes at 4°C, and the supernatant was collected. ELISA kits from Meimian Biology Company in China were used to measure the NE content in the serum, kidney of six rats in each group, following the instructions outlined in the kit manual.

### 2.6 Bioinformatics Analysis

Principal Component Analysis (PCA) was performed using theR package gmodels (https://www.r-project.org/). Differentiallyexpressed genes (DEGs), Kyoto Encyclopedia of Genes andGenomes (KEGG) pathway analysis, Protein–Protein Interaction(PPI) networks and Gene Set Enrichment Analysis. The authors declare that all supporting data are available within the article and its online supplementary files.

### 2.7 Statistical Analyses

All data are presented as mean ± standard error (SEM). Statistical analyses were performed using SPSS 25.0 software. Comparisons among multiple groups were conducted using one-way ANOVA followed by Tukey’s post hoc test or Fisher’s least significant difference test. A value of p < 0.05 was considered statistically significant.

## 3 RESULTS

### 3.1 RDN Reduced Blood Pressure in SHRs

Sham-operated SHRs exhibited sustained and severe hypertension. RDN treatment significantly reduced systolic blood pressure to a similar degree as Sac/Val. No significant difference in antihypertensive efficacy was observed between RDN and Sac/Val groups.

**Figure 1.**
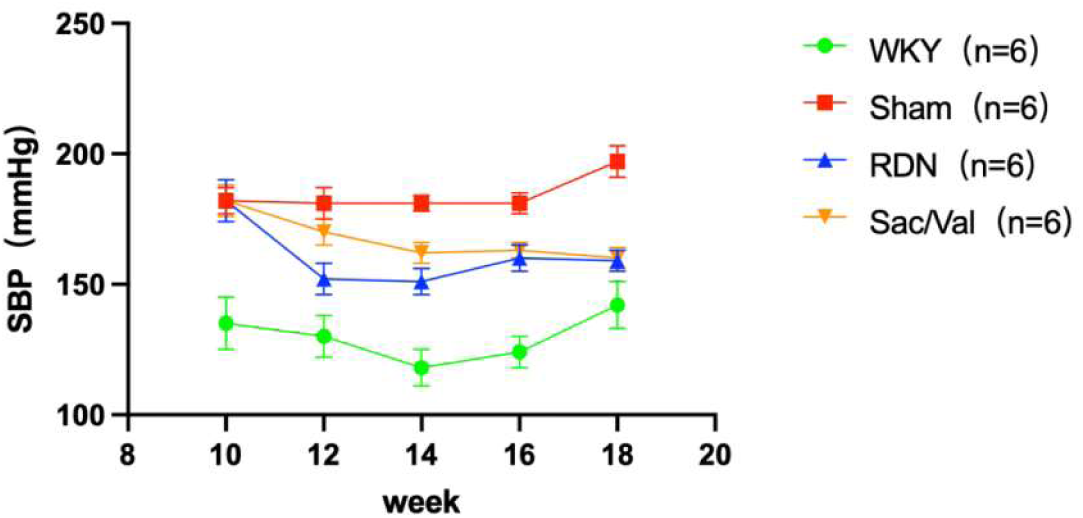
Time course of systolic blood pressure (SBP) in rats following different interventions. Systolic blood pressure (SBP) was measured by the tail-cuff method in four groups: normotensive Wistar-Kyoto (WKY) rats (green circles, n=6), spontaneously hypertensive rats (SHR) with sham surgery (red squares, n=6), SHR treated with renal denervation (RDN, blue triangles, n=6), and SHR treated with sacubitril/valsartan (Sac/Val, orange triangles, n=6). SBP was recorded at baseline (10 weeks of age) and every 2 weeks thereafter until week 18. The Sham group showed progressive and severe hypertension, while both RDN and Sac/Val significantly reduced SBP compared with Sham, with no significant difference in antihypertensive efficacy between the two treatment groups. Data are presented as mean ± standard error (SEM). Repeated-measures two-way ANOVA was performed for statistical analysis.

### 3.2 RDN Inhibited Renal Sympathetic Nervous System Activity

Plasma norepinephrine (NE) and renal tyrosine hydroxylase (TH) expression were measured as markers of sympathetic nervous system activity. Compared with the WKY group, the Sham-operated SHRs exhibited significantly elevated plasma NE concentrations (****p< 0.0001 vs WKY), consistent with sympathetic overactivation. RDN treatment significantly reduced plasma NE levels to values comparable to those observed in the Sac/Val group (both ****p < 0.0001 vs Sham), with no significant difference between the two interventions.

Consistent with plasma NE data, Western blot analysis showed markedly increased TH protein expression in the renal arteries of Sham SHRs. RDN significantly downregulated renal TH expression compared with the Sham group (***p < 0.001), confirming effective inhibition of renal sympathetic nerve activity.

These findings indicate that RDN exerts a potent sympatholytic effect in SHRs, which likely contributes to its antihypertensive and vascular protective effects.

**Figure 3.2.1.**
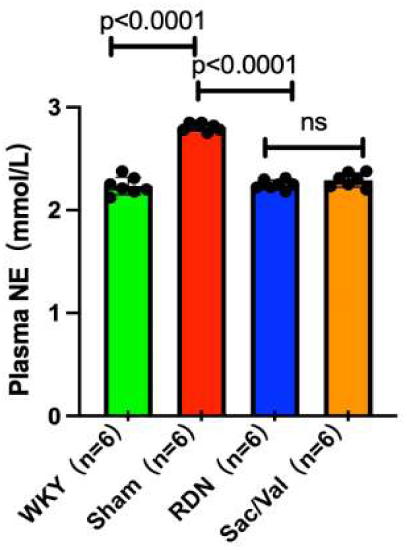
Effects of renal denervation (RDN) and sacubitril/valsartan (Sac/Val) on plasma norepinephrine (NE) levels in spontaneously hypertensive rats (SHRs). Plasma NE concentrations were measured in four groups: Wistar-Kyoto (WKY) rats, sham-operated SHRs, RDN-treated SHRs, and Sac/Val-treated SHRs (n=6 per group). Plasma NE levels were significantly higher in the Sham group compared with the WKY group (****p < 0.0001). Both RDN and Sac/Val interventions significantly reduced plasma NE levels compared with the Sham group (****p < 0.0001). No significant difference was observed between the RDN and Sac/Val groups (ns, not significant), indicating that both interventions effectively suppress sympathetic nervous system activity in SHRs. Data are presented as mean ± standard error (SEM). One-way ANOVA followed by Tukey’s post hoc test was used for statistical analysis.

**Figure 3.2.2.**
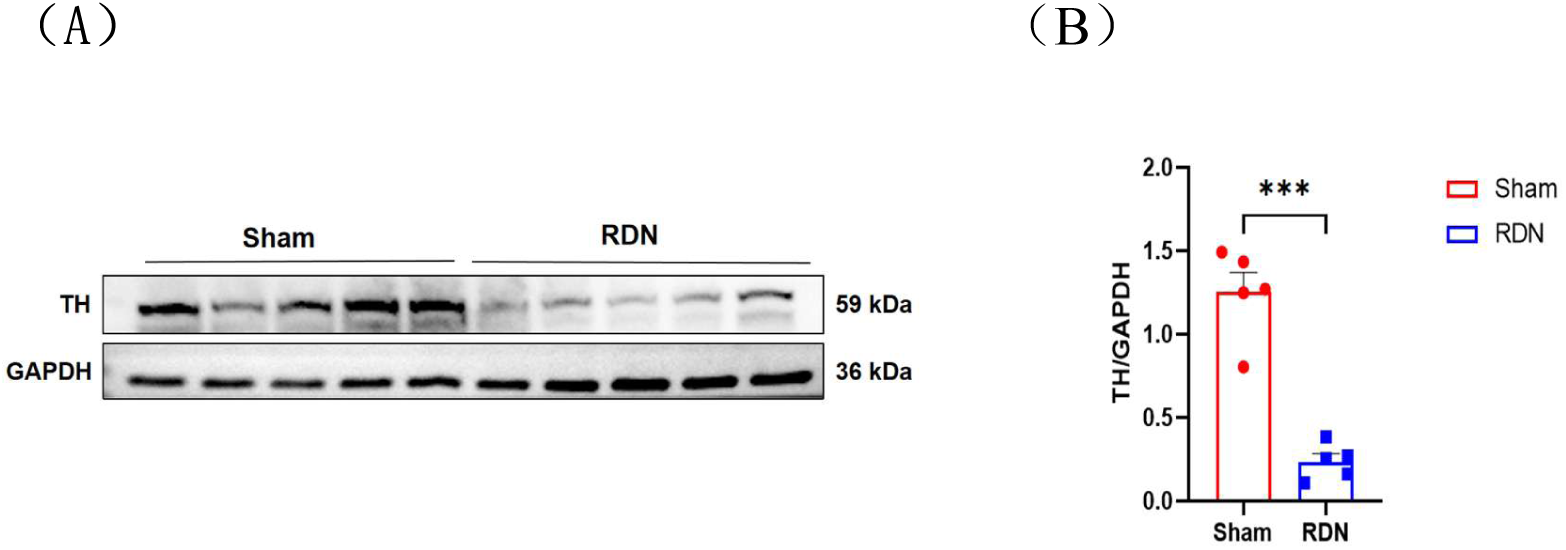
Effects of renal denervation (RDN) on renal artery tyrosine hydroxylase (TH) protein expression in spontaneously hypertensive rats (SHRs). (A) Representative Western blot bands showing TH (59 kDa) protein expression in renal arteries from the Sham and RDN groups. GAPDH (36 kDa) was used as an internal control. (B) Quantitative analysis of TH protein expression normalized to GAPDH. Compared with the Sham group, RDN significantly reduced TH protein expression in the renal artery (***p < 0.001). These results indicate that RDN effectively inhibits renal sympathetic nerve activity in SHRs. Data are presented as mean ± standard error (SEM). Student’s t-test was used for statistical analysis.

### 3.3 RDN Attenuated Mesenteric Artery Remodeling

Sham SHRs displayed severe vascular remodeling characterized by increased wall-to-lumen area ratio, wall thickness-to-lumen radius ratio, and collagen deposition. Both RDN and Sac/Val significantly improved these structural abnormalities, with comparable therapeutic effects.

**Figure 3.3.1.**
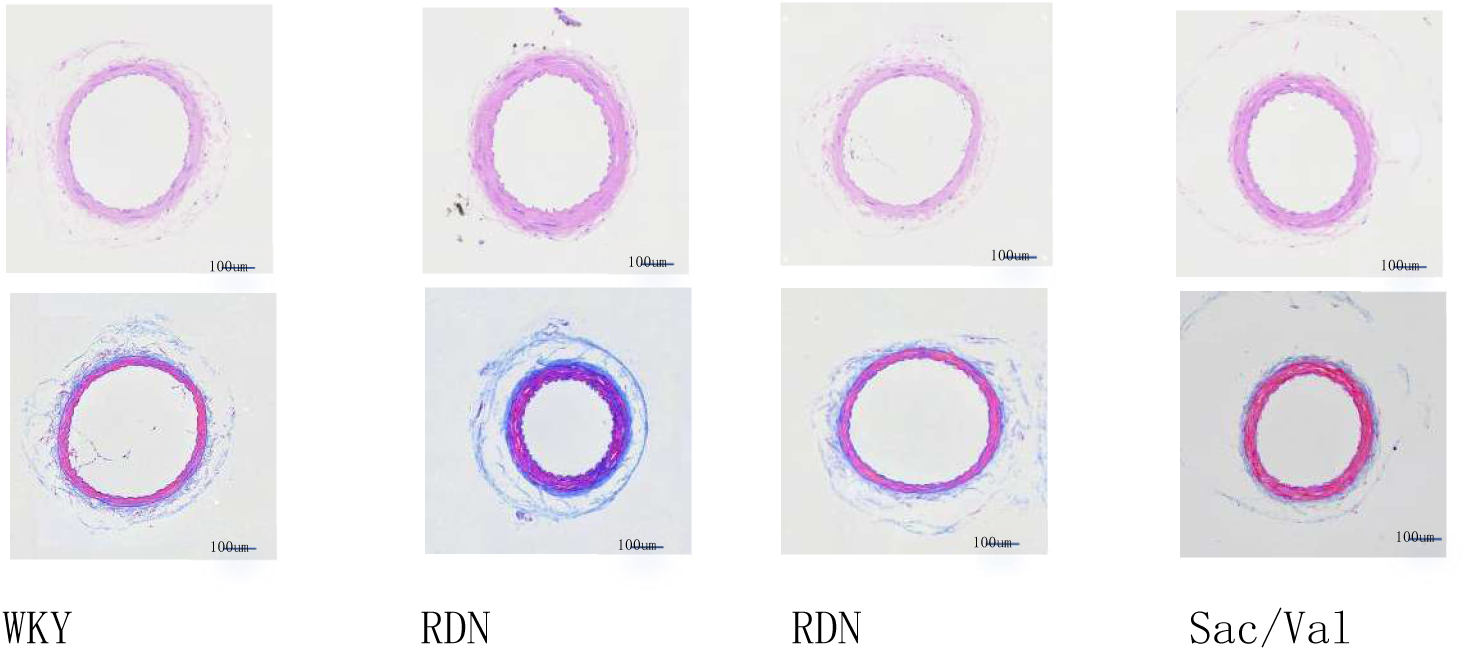
Effects of renal denervation (RDN) and sacubitril/valsartan (Sac/Val) on mesenteric artery remodeling in spontaneously hypertensive rats (SHRs). Representative images of hematoxylin-eosin (HE) staining (upper panel) and Masson’s trichrome staining (lower panel) of mesenteric arteries from Wistar-Kyoto (WKY) rats, Sham-operated SHRs, RDN-treated SHRs, and Sac/Val-treated SHRs (scale bar = 100 μm). Sham-operated SHRs exhibited severe vascular remodeling, characterized by increased wall-to-lumen ratio, disorganized smooth muscle cell arrangement, and excessive collagen deposition. Both RDN and Sac/Val significantly attenuated these structural abnormalities, with reduced wall thickening and improved vascular architecture.

One-way ANOVA followed by Tukey’s post hoc test was used to compare the wall-to-lumen area ratios among the four groups at 14 and 18 weeks of age. At 14 weeks of age, there was a significant overall difference among groups (F=4.78, p=0.012). Post hoc tests showed that the ratios were significantly higher in the Sham (p=0.022) and Sac/Val (p=0.016) groups compared with the WKY group. At 18 weeks of age, the overall difference remained significant (F=14.35, p<0.001). The Sham group exhibited a significantly higher ratio than the WKY group (p<0.001), while both Sac/Val (p=0.001) and RDN (p<0.001) treatment significantly reduced the ratio compared with the Sham group. No significant differences were observed between the Sac/Val and RDN groups at either time point.

**Figure 3.3.2.**
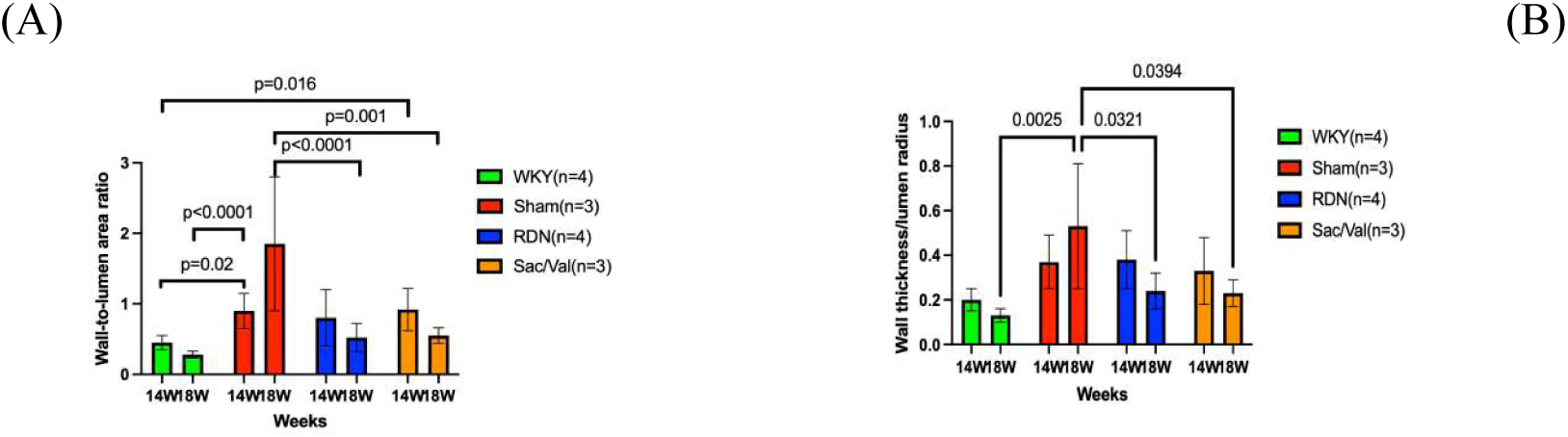
Effects of renal denervation (RDN) and sacubitril/valsartan (Sac/Val) on mesenteric artery remodeling in spontaneously hypertensive rats (SHRs). (A) Wall-to-lumen area ratio (W/L) of mesenteric arteries at 14 and 18 weeks of age (equal 4 and 8 weeks post-intervention). (B) Wall thickness-to-lumen radius ratio (WT/LR) of mesenteric arteries at 14 and 18 weeks of age. Compared with the WKY group, the Sham group exhibited significantly increased W/L and WT/LR ratios, indicating severe vascular remodeling. Both RDN and Sac/Val significantly reduced these structural indices at both time points compared with the Sham group, demonstrating comparable therapeutic efficacy. Data are presented as mean ± SEM. One-way ANOVA followed by Tukey’s post hoc test was used for statistical analysis.

### 3.4 RDN Restored Endothelial and Vascular Smooth Muscle Function

To evaluate endothelial function, acetylcholine (ACh)-induced endothelium-dependent relaxation was measured in isolated mesenteric arteries.

Compared with the WKY group, the Sham group showed a marked rightward shift of the concentration-response curve and a significant reduction in maximum relaxation (Emax), accompanied by a significantly decreased pD₂ value (all p < 0.001). Both Sac/Val and RDN treatments significantly restored ACh-induced relaxation, with a leftward shift of the curve and increased Emax and pD₂ values compared with the Sham group (all p < 0.001). The pD₂ value in the RDN group was comparable to that in the WKY group (p > 0.05), whereas the Sac/Val group still exhibited a significant difference (p < 0.01).

To further assess vascular smooth muscle function, sodium nitroprusside (SNP)-induced endothelium-independent relaxation was determined. The Sham group displayed a pronounced rightward shift of the SNP concentration-response curve and reduced Emax, indicating impaired smooth muscle relaxation (p < 0.001 vs WKY).

Both Sac/Val and RDN interventions significantly improved SNP-induced relaxation, as evidenced by the leftward shift of the curve and elevated Emax and pD₂ values (all p < 0.001 vs Sham). The pD₂ values in both treated groups remained partially different from the WKY group (p < 0.01 for Sac/Val, p< 0.05 for RDN). No significant difference was observed between the Sac/Val and RDN groups in either ACh- or SNP-induced relaxation (all p> 0.05).

**Figure 3.4.**
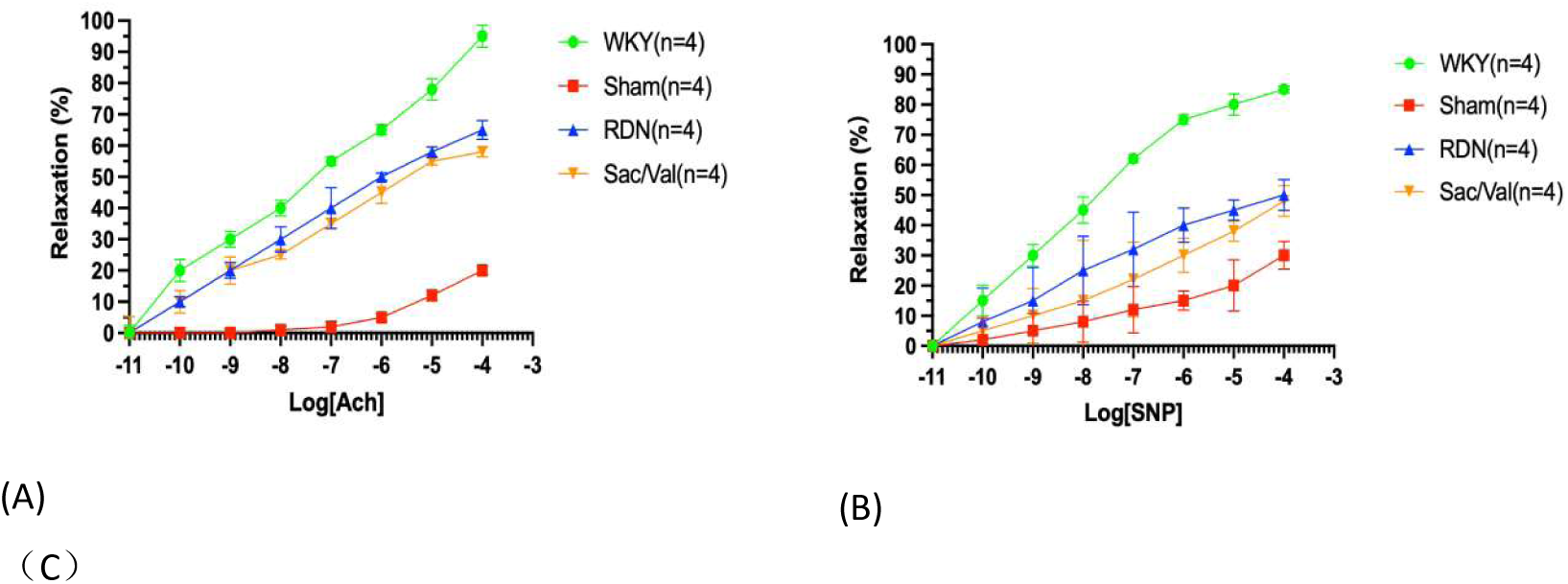
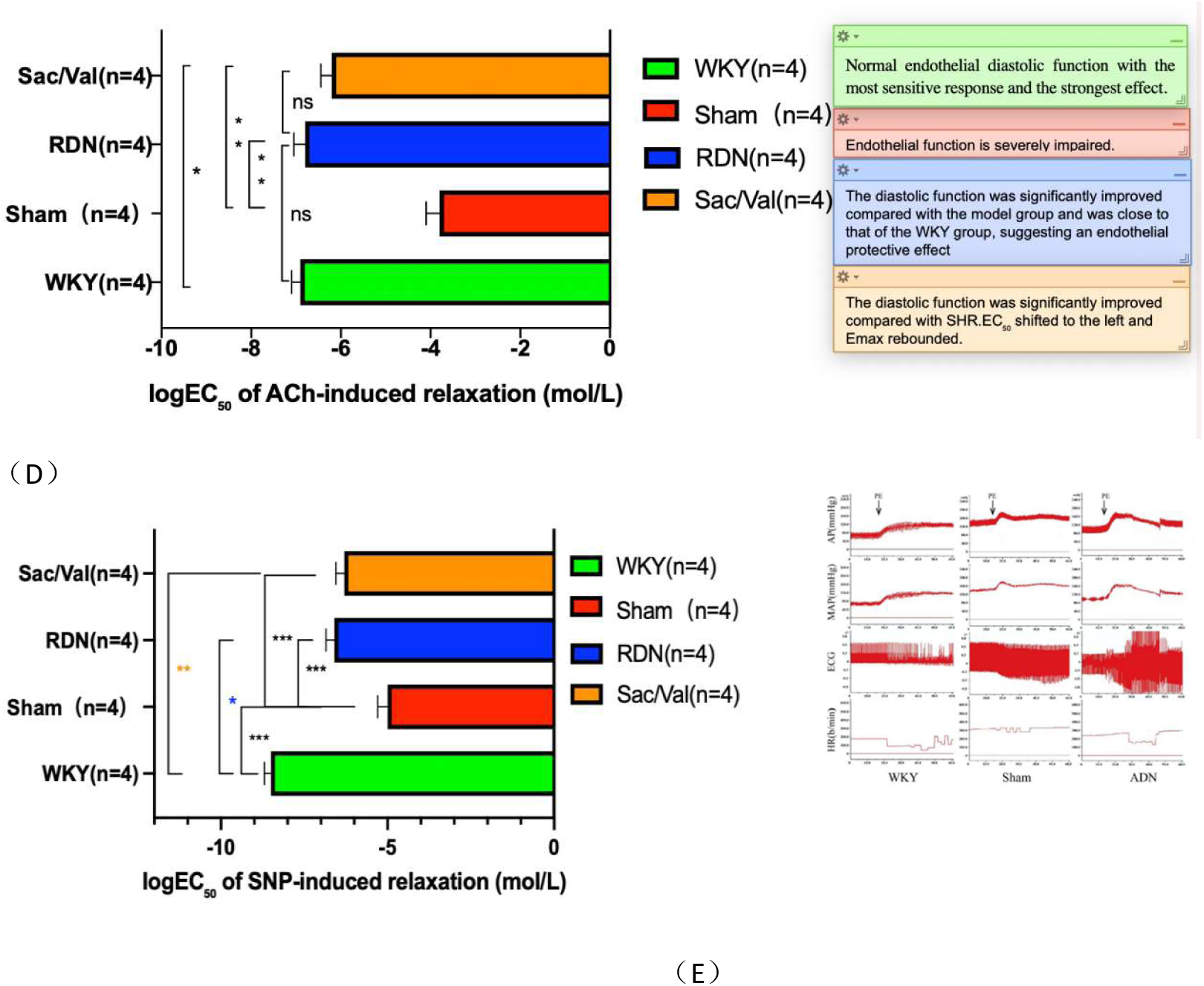
Endothelium-dependent and endothelium-independent relaxation in mesenteric arteries. (A): Acetylcholine (ACh)-induced endothelium-dependent relaxation in isolated mesenteric arteries from WKY, Sham, Sac/Val, and RDN groups. Vessels were pre-contracted with phenylephrine, and relaxation responses to cumulative concentrations of ACh (10⁻¹¹ to 10⁻⁴ mol/L) were expressed as a percentage of the pre-contraction level. Concentration-response curves were fitted by non-linear regression using a sigmoidal dose-response model with variable slope. Compared with the WKY group, the Sham group showed a marked rightward shift and significantly reduced maximum relaxation (Emax), indicating severe endothelial dysfunction. Both Sac/Val and RDN significantly restored relaxation responses, with increased Emax and leftward-shifted curves toward the WKY group. (B): Sodium nitroprusside (SNP)-induced endothelium-independent relaxation in the same groups. Relaxation was expressed as a percentage of phenylephrine-induced precontraction. Curves were fitted using a non-linear regression model (log[agonist] vs. response, variable slope).Data are presented as mean ± SEM. (C)(D): One-way ANOVA followed by Tukey’s multiple comparisons test revealed a significant difference in logEC₅₀ among groups (P Compared with the WKY group, the Sham group exhibited a marked rightward shift of logEC₅₀ (p< 0.001) indicating severely impaired endothelial sensitivity. Both Sac/Val and RDN treatments significantly shifted logEC₅₀ leftward compared with the Sham group (both p< 0.001). The logEC₅₀ value in the RDN group was comparable to that in the WKY group (p > 0.05) whereas a significant difference remained between the Sac/Val and WKY groups (p < 0.01). There was no significant difference between the RDN and Sac/Val groups (p > 0.05). (E): Physiological curve record graphs, representative change curves of arterial pressure (AP), mean arterial pressure (MAP), electrocardiogram (ECG), and heart rate (HR) in each group of rats (WKY, Sham, RDN) after intravenous injection of phenylephrine (PE), used to evaluate the arterial pressure reflex response.

### 3.5 RDN Improved Arterial Baroreflex Sensitivity

Arterial baroreflex sensitivity (BRS) was markedly reduced in spontaneously hypertensive rats (SHRs). A two-way repeated-measures ANOVA revealed a significant main effect of group (p < 0.001) and a significant group × time interaction (P = 0.010), while the main effect of time was not statistically significant (P = 0.084). At 18 weeks of age, BRS was significantly lower in the Sham group compared with the normotensive WKY control group (p < 0.001). Both renal denervation (RDN) and sacubitril/valsartan (Sac/Val) significantly improved BRS relative to the Sham group (p = 0.021 for both interventions). At 18 weeks of age, BRS remained severely depressed in the Sham group (p < 0.001 vs WKY). The beneficial effect of Sac/Val on BRS persisted at 18 weeks of age (p < 0.001 vs Sham), whereas the improvement in the RDN group was no longer statistically significant (p = 0.186 vs Sham). Notably, BRS in the Sac/Val group was significantly higher than in the RDN group at 18 weeks of age (p = 0.004).

Correlation analysis further demonstrated that the expression level of the TLR4/NF-κB pathway in mesenteric arteries was negatively correlated with BRS (r = −0.78, p < 0.01) and positively correlated with systolic blood pressure (r = 0.82, p < 0.01).

These findings suggest that both RDN and Sac/Val improve baroreflex function in SHRs at early time points, with the effect of Sac/Val being more sustained. The inverse relationship between TLR4/NF-κB activity and BRS supports the notion that vascular inflammation contributes to impaired baroreflex regulation in hypertension.

**Figure 3.5.1.**
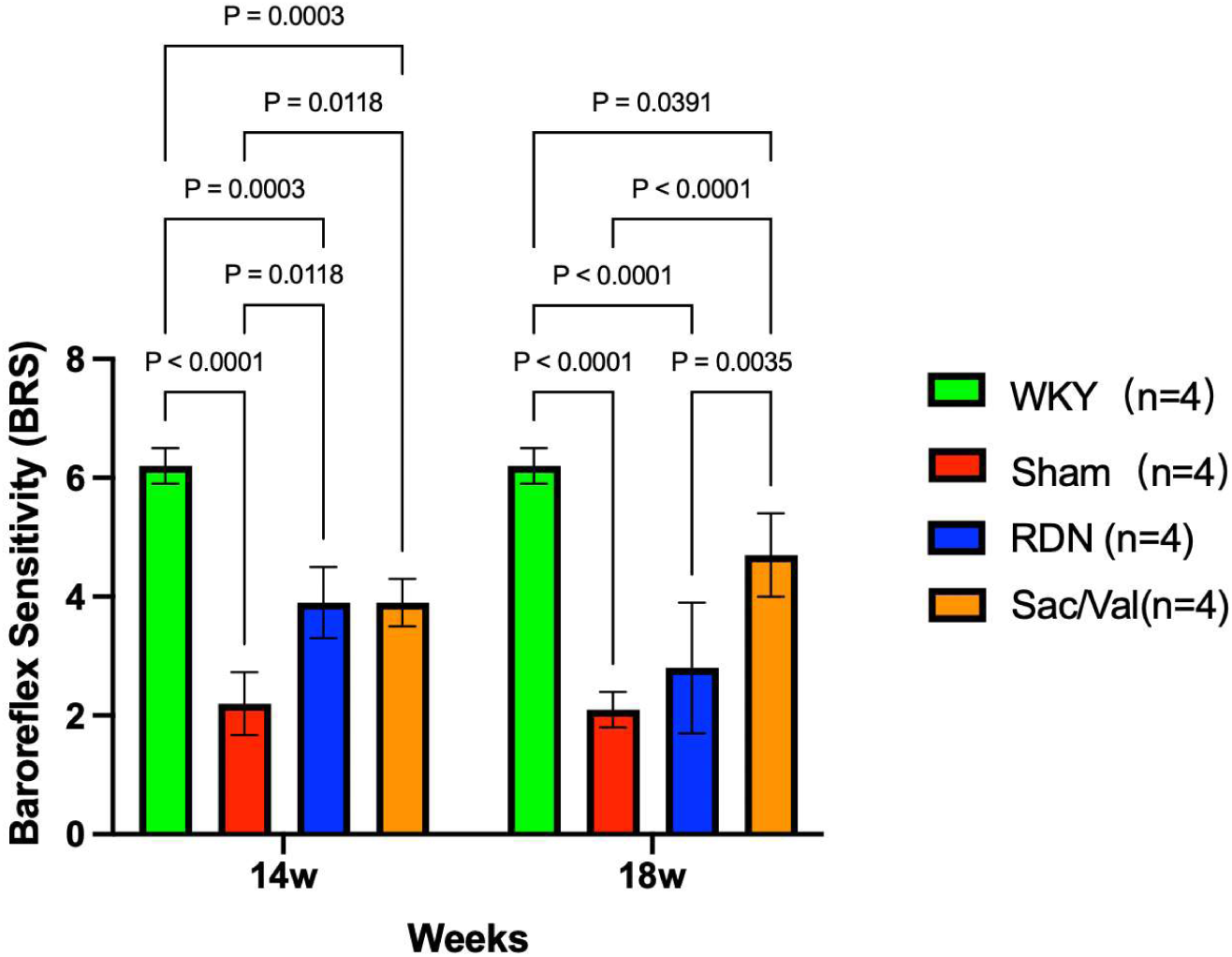
Forest plot of baroreflex sensitivity (BRS) in WKY, SHAM, Sac/Val, and RDN groups at 14 and 18 weeks of age. Data are presented as mean with 95% confidence intervals. BRS was significantly reduced in the SHAM group. Sac/Val treatment improved BRS in a time-dependent manner, while the effect of RDN was not sustained at 18 weeks of age.

### 3.6 Verify the changes of miR128 after RDN intervention and identify the mediators mediating the effect of mir-128 on vascular inflammation in SHR

3.6.1 RDN Downregulates miR-128 Expression in the Mesenteric Arteries of SHRs Given the high sequence homology of miR-128 between humans and rats (Figure 3.6.1.), we assessed miR-128 expression in the mesenteric arteries of SHRs. One-way ANOVA revealed a significant overall difference among the four groups (F = 45.69, p < 0.0001). Post hoc Tukey’s test showed that miR-128 expression was significantly upregulated in the Sham-operated SHRs compared with the normotensive WKY group (p < 0.0001). Renal denervation (RDN) significantly reduced miR-128 levels compared with both the Sham group and the Sac/Val group (p < 0.0001 for both). Notably, no significant difference in miR-128 expression was observed between the Sham and Sac/Val groups (p = 0.9123). These findings indicate that miR-128 is selectively downregulated by RDN, independent of blood pressure reduction.

**Figure 3.6.1.1.**
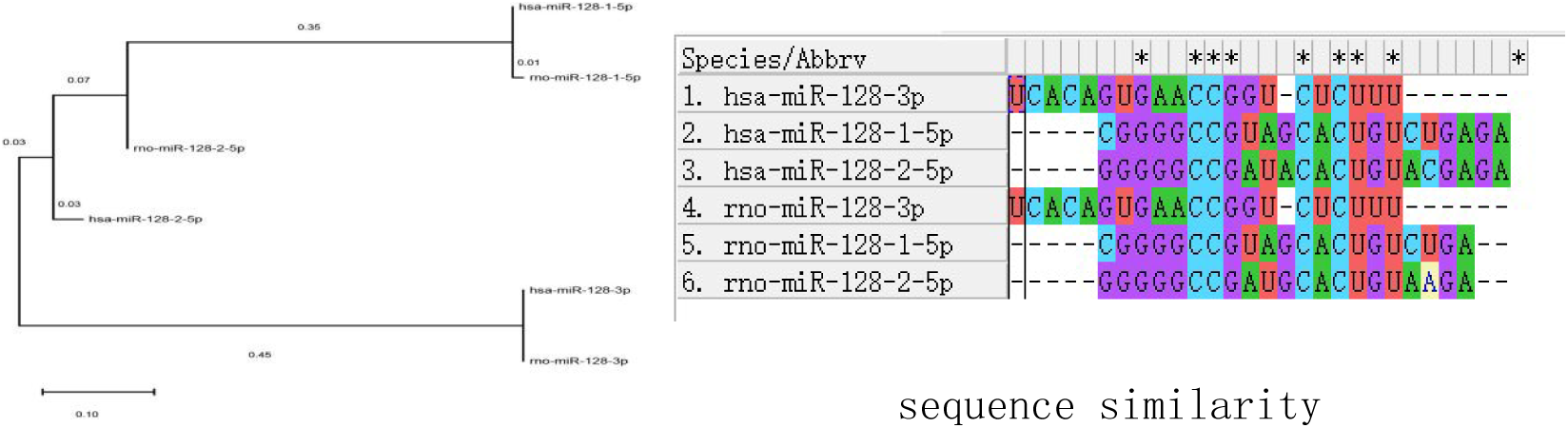
Sequence homology analysis of miR-128 between humans and rats. The Neighbor-Joining phylogenetic tree (left panel) and multiple sequence alignment (right panel) demonstrate high sequence conservation of miR-128-3p between human (hsa) and rat (rno) orthologs. The seed region, which is critical for target recognition, is highly conserved across species, supporting the translatability of findings from rat models to human studies.

**Figure 3.6.1.2.**
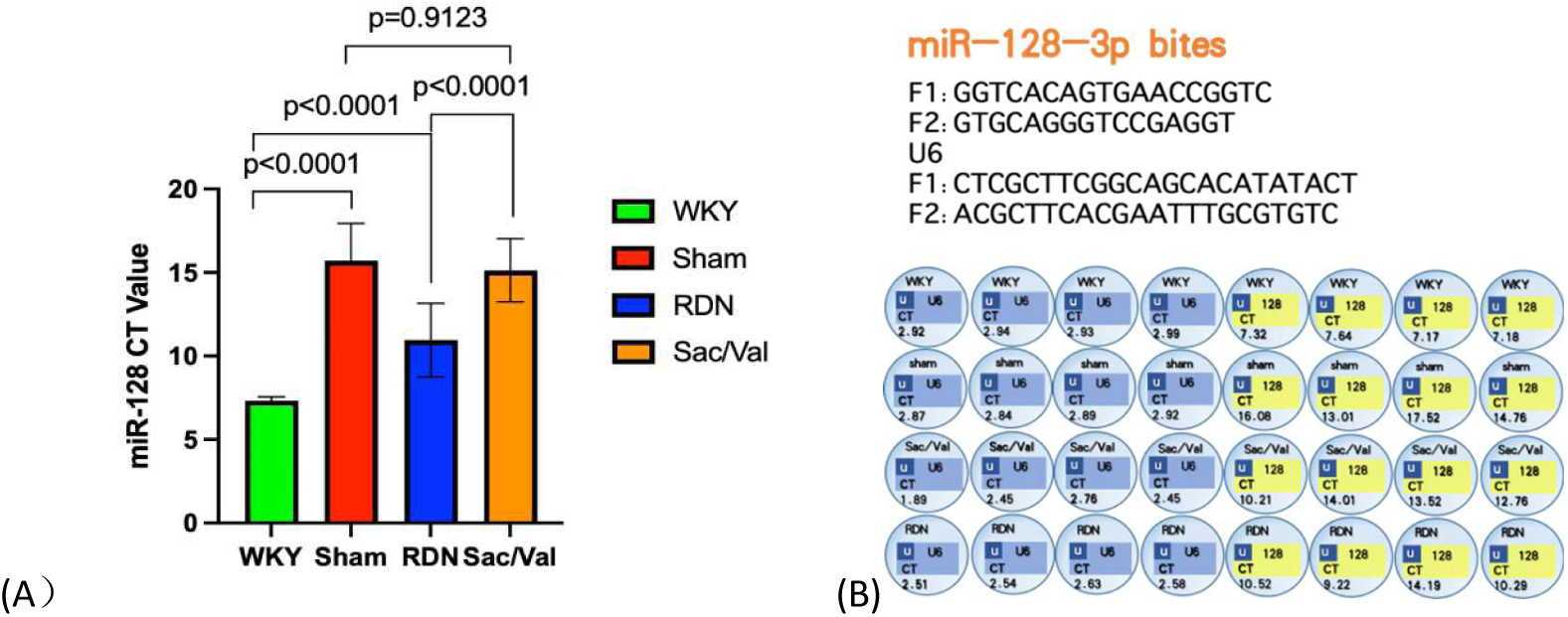
Regulation of miR-128 expression by renal denervation (RDN) in spontaneously hypertensive rats (SHRs). (A) Heatmap showing the expression levels of miR-128-3p in mesenteric arteries of rats from the WKY, Sham, RDN, and Sac/Val groups. (B) Quantitative analysis of miR-128 expression (CT values) in mesenteric arteries. Compared with the WKY group, miR-128 expression was significantly upregulated in the Sham group (p < 0.0001). Compared with the Sham group, miR-128 expression was significantly reduced in the RDN group (p < 0.0001), while no significant difference was observed in the Sac/Val group (p = 0.9123 vs Sham). These results indicate that RDN specifically suppresses miR-128 expression in hypertensive rats, independent of blood pressure reduction. Data are presented as mean ± SEM. One-way ANOVA followed by Tukey’s post hoc test was used for statistical analysis.

#### 3.6.2 RDN Inhibits Activation of the TLR4/NF-κB Signaling Pathway in Mesenteric Arteries of SHRs

Compared with the normotensive WKY rats, SHRs in the Sham group exhibited significantly elevated expression of TLR4 in the mesenteric arteries. Concurrently, the phosphorylation of NF-κB p65, as well as the expression levels of pro-inflammatory cytokines TNF-α IL-6 and IL-1β, were markedly increased in Sham-operated SHRs. Renal denervation (RDN) significantly suppressed the activation of the TLR4/NF-κB pathway, as evidenced by reduced TLR4 expression, decreased NF-κB p65 phosphorylation, and downregulated production of TNF-α IL-6 and IL-1β (Figure 1.2.2). These findings suggest that activation of the TLR4/NF-κB inflammatory cascade promotes vascular inflammation and exacerbates pathological vascular remodeling in hypertension. RDN may improve vascular function and regulate blood pressure, at least in part, by inhibiting NF-κB-mediated vascular inflammation, providing experimental evidence for the anti-inflammatory mechanism underlying the vascular protective effects of RDN.

**Figure 3.6.2.1.**
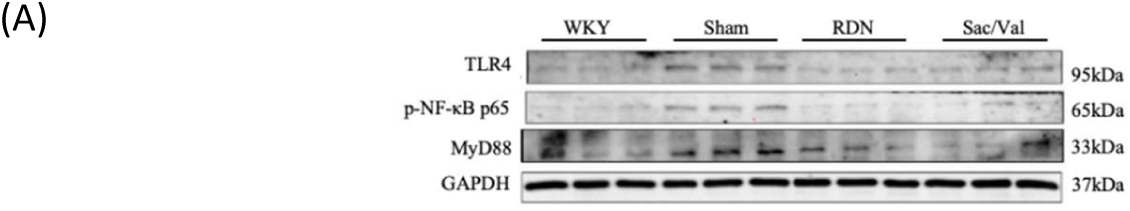
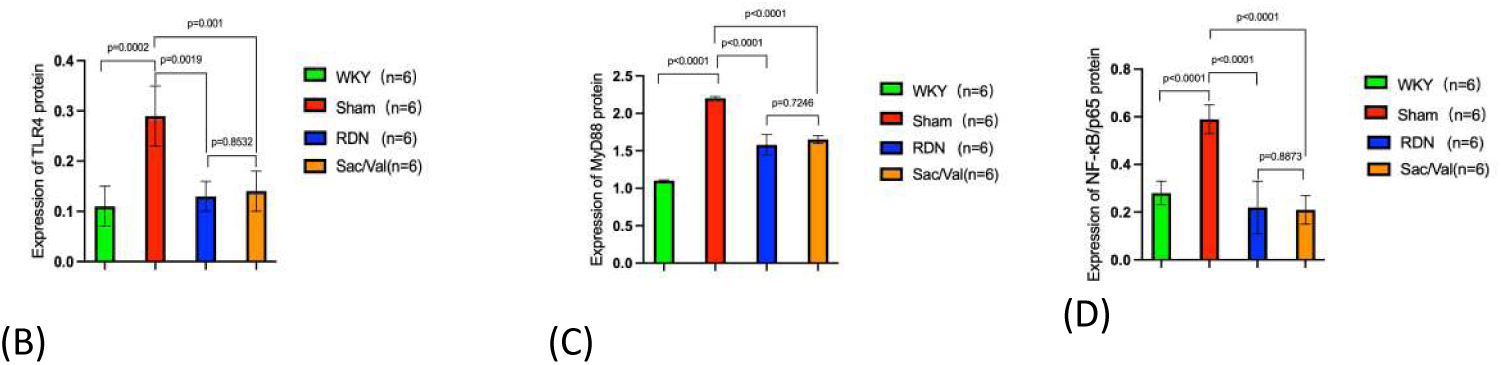
Effects of renal denervation (RDN) on the TLR4/NF-κB signaling pathway in mesenteric arteries of spontaneously hypertensive rats (SHRs). (A) Representative Western blot bands showing the protein expression of TLR4 (95 kDa), phosphorylated NF-κB p65 (p-NF-κB p65, 65 kDa), and MyD88 (33 kDa) in mesenteric arteries from Wistar-Kyoto (WKY), Sham, RDN, and sacubitril/valsartan (Sac/Val) groups. GAPDH (37 kDa) was used as the internal control. (B-D) Quantitative analysis of the relative protein expression levels of TLR4, MyD88, and p-NF-κB p65, respectively, normalized to GAPDH. Compared with the WKY group, the Sham group exhibited significantly upregulated expression of TLR4, MyD88, and p-NF-κB p65. Both RDN and Sac/Val significantly reduced the expression of these inflammatory signaling proteins compared with the Sham group. Data are presented as mean ± SEM (n=6 per group). One-way ANOVA followed by Tukey’s post hoc test was used for statistical analysis.

**Figure 3.6.2.2.**
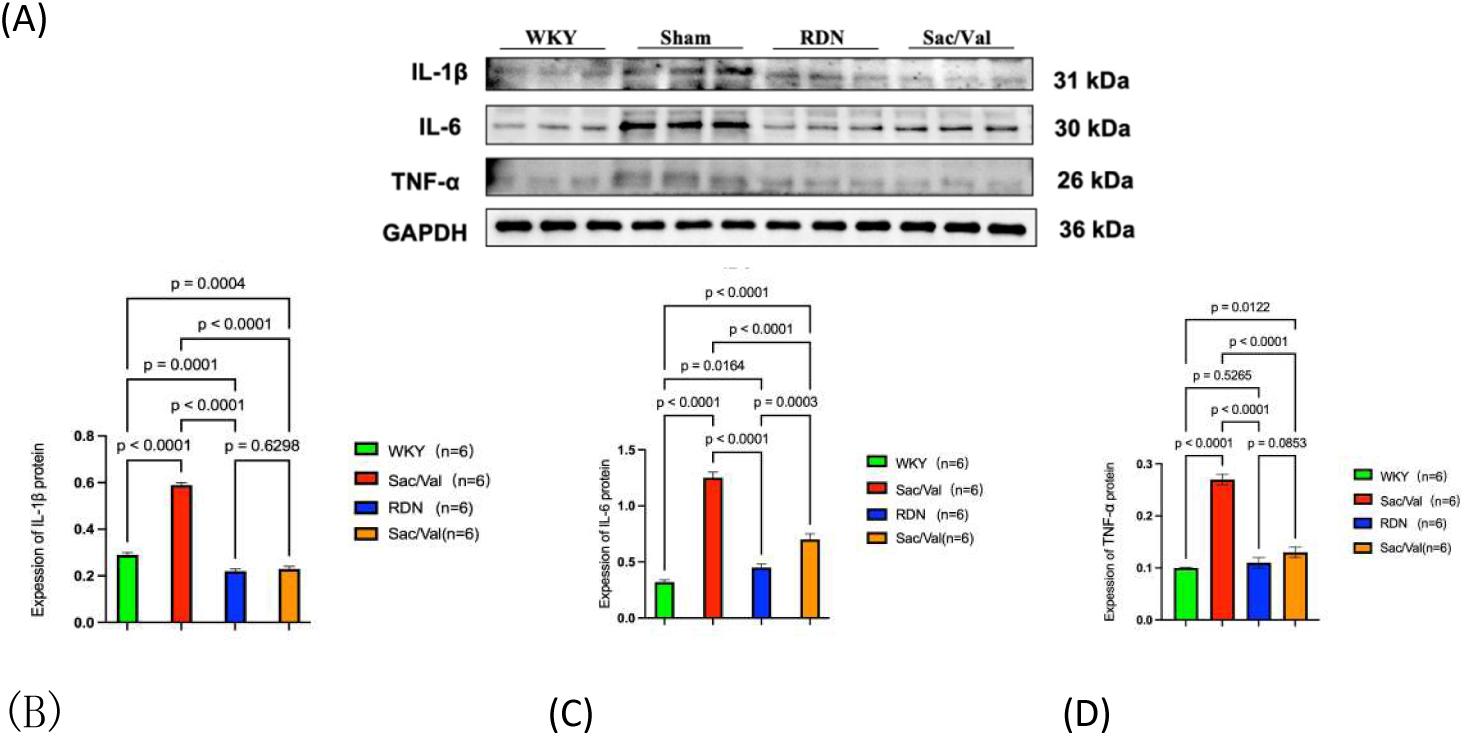
Effects of renal denervation (RDN) on pro-inflammatory cytokine expression in mesenteric arteries of spontaneously hypertensive rats (SHRs). (A) Representative Western blot bands showing the protein expression of interleukin-1β (IL-1β, 31 kDa), interleukin-6 (IL-6, 30 kDa), and tumor necrosis factor-α (TNF-α, 26 kDa) in mesenteric arteries from Wistar-Kyoto (WKY), Sham, RDN, and sacubitril/valsartan (Sac/Val) groups. GAPDH (36 kDa) was used as the internal control. (B-D) Quantitative analysis of the relative protein expression levels of IL-1β, IL-6, and TNF-α, respectively, normalized to GAPDH. Compared with the WKY group, the Sham group exhibited significantly upregulated expression of all three pro-inflammatory cytokines. Both RDN and Sac/Val significantly reduced cytokine expression compared with the Sham group. Data are presented as mean ± SEM (n=6 per group). One-way ANOVA followed by Tukey’s post hoc test was used for statistical analysis.

### 3.7 miR**-**128 Directly Targets PPAR**-γ**

Transcriptomic analysis and bioinformatic prediction identified PPAR-γ as a key target gene of miR-128 within the PPAR signaling pathway. PPAR-γ expression was significantly decreased in Sham SHRs and was restored after RDN.

Our research team conducted transcriptome sequencing on the mesenteric artery tissues of SHR patients who underwent sham surgery and SHR patients who received RDN treatment. Use the absolute value of logfc >1 and padj<0.05 as the threshold to screen for differential genes.

Compared with the sham group, the RDN group obtained 640 up-regulated genes and 169 down-regulated genes. A total of 581 regulated mrnas of rat miR128 were predicted using an online website. Subsequently, the differentially expressed genes in the transcriptome were interchanged with the genes regulated by miR128, resulting in a total of 29 genes. Through DAVID’s GO and KEGG analyses, it was found that the PPAR signaling pathway was a significantly enriched pathway (Figures 3.7.1. and 3.7.2.). Analysis of abnormal genes in the PPAR signaling pathway and miR128-2-3P targets in SHR after RDN revealed key pathways. PPAR-γ in the pathway was identified as a key regulatory target (Table 1).

**Figure 3.7.1.**
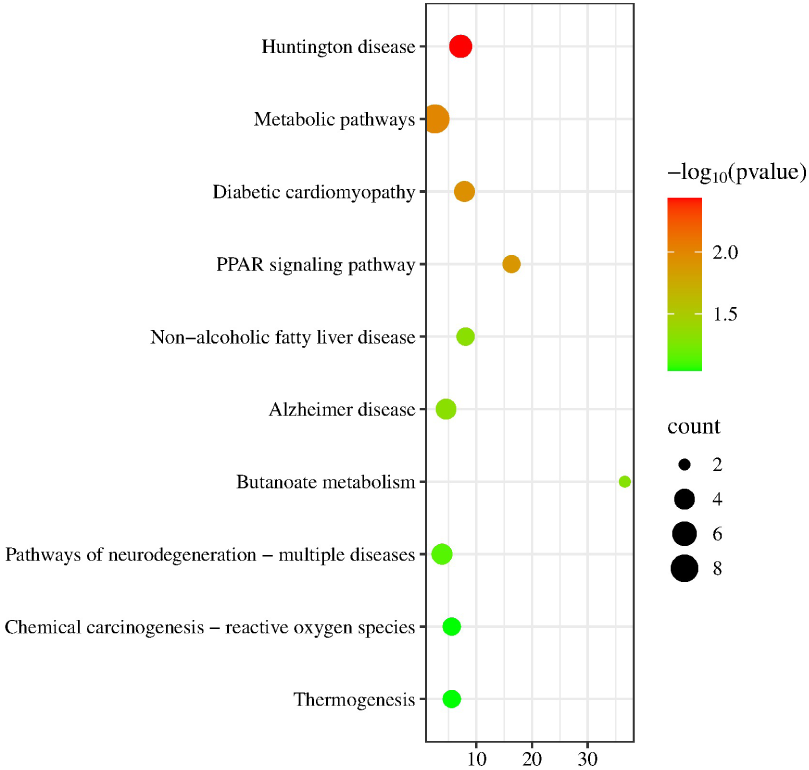
Bubble plot of enriched signaling pathways for differentially expressed genes.

**Figure 3.7.2.**
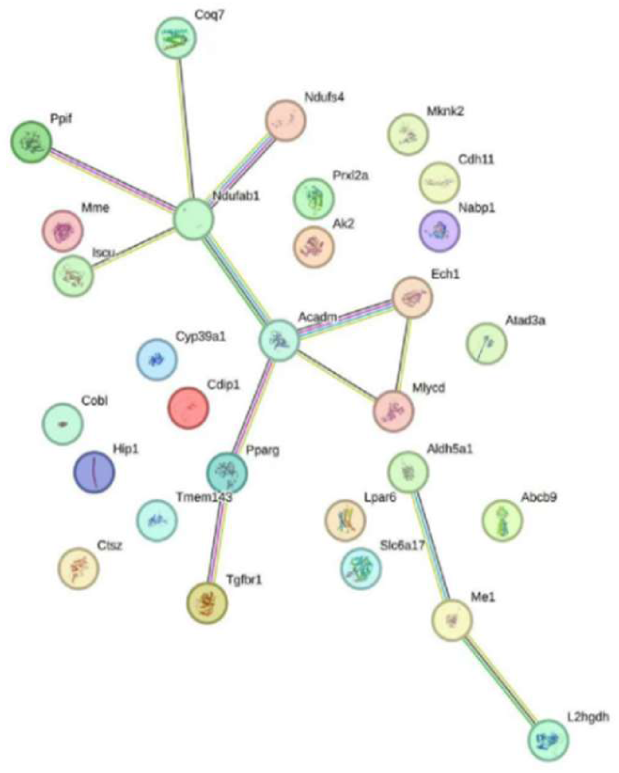
Protein–protein interaction (PPI) network of key targets

**Table 1.** Transcriptomic identification of PPAR-γ and its association with the TLR4/NF-κB inflammatory pathway.

| Category | Term | Genes | Count | List Total | Pop Hits | Pop Total | P-Value | Fisher Exact |
| --- | --- | --- | --- | --- | --- | --- | --- | --- |
| KEGG_PATHWAY | Huntington disease | 17% | 5 | 20 | 344 | 9898 | 0.00367 | 0.000496 |
| KEGG_PATHWAY | Metabolic pathways | 31% | 9 | 20 | 1724 | 9898 | 0.0102 | 0.00384 |
| KEGG_PATHWAY | Diabetic cardiomyopathy | 14% | 4 | 20 | 252 | 9898 | 0.0117 | 0.00144 |
| KEGG_PATHWAY | PPAR signaling pathway | 10% | 3 | 20 | 91 | 9898 | 0.0129 | 0.000765 |
| KEGG_PATHWAY | Non-alcoholic fatty liver disease | 10% | 3 | 20 | 184 | 9898 | 0.0478 | 0.00571 |
| KEGG_PATHWAY | Alzheimer disease | 14% | 4 | 20 | 434 | 9898 | 0.0481 | 0.0101 |
| KEGG_PATHWAY | Butanoate metabolism | 7% | 2 | 20 | 27 | 9898 | 0.0506 | 0.00132 |
| KEGG_PATHWAY | Pathways of neurodegeneration - multiple diseases | 14% | 4 | 20 | 515 | 9898 | 0.073 | 0.018 |
| KEGG_PATHWAY | Chemical carcinogenesis - reactive oxygen species | 10% | 3 | 20 | 266 | 9898 | 0.0912 | 0.0156 |
| KEGG_PATHWAY | Thermogenesis | 10% | 3 | 20 | 266 | 9898 | 0.0912 | 0.0156 |

## 4 DISCUSSION

Hypertension-induced vascular remodeling is a key driver of cardiovascular injury and target organ damage[15,16]. The present study demonstrates that RDN exerts potent vascular protective effects in SHRs beyond blood pressure control[17,18]. We identify a novel molecular axis—miR-128/PPAR-γ/TLR4/NF-κB—that mediates the beneficial effects of RDN on vascular remodeling and inflammation[2].

First, we confirmed that RDN effectively reduces blood pressure and inhibits renal sympathetic overactivity in SHRs, with efficacy comparable to Sac/Val. Beyond hemodynamic improvement, RDN significantly attenuated structural remodeling, restored endothelial and vascular smooth muscle function, and improved arterial baroreflex sensitivity[5]. These findings support the concept that RDN provides direct vascular protection independent of blood pressure reduction[6,19].

Second, we identified miR-128 as a key molecular target specific to RDN. MiR-128 was upregulated in hypertensive arteries and was selectively suppressed by RDN, but not by Sac/Val, indicating that miR-128 modulation is a distinct molecular response to RDN rather than a general effect of blood pressure lowering[8,12].

Third, we demonstrated that miR-128 directly targets PPAR-γ. PPAR-γ expression was reduced in SHRs and was restored by RDN[11]. Upregulation of PPAR-γ subsequently inhibited the TLR4/NF-κB inflammatory pathway and reduced the production of pro-inflammatory cytokines. These results establish a mechanistic cascade: RDN → ↓miR-128 → ↑PPAR-γ → inhibition of TLR4/NF-κB → ↓vascular inflammation → improved vascular remodeling.

This study extends previous observations by linking non-coding RNA regulation to the anti-inflammatory and vascular-protective effects of RDN[11]. The miR-128/PPAR-γ/TLR4/NF-κB axis represents a novel and promising therapeutic target for hypertensive vascular remodeling[13].

Several limitations should be acknowledged. First, although the present study was conducted in an animal model, we are currently performing in vitro mechanistic validation experiments. Specifically, we are conducting vascular smooth muscle cell (VSMC) culture assays to further confirm the direct binding relationship between miR-128 and PPAR-γ at the cellular level. Dual-luciferase reporter assays and gain-and loss-of-function studies in VSMCs are underway to conclusively verify the targeted regulation of PPAR-γ by miR-128. These in vitro experiments will provide solid cellular evidence to support the molecular cascade identified in vivo. Second, despite the confirmation of miR-128 dysregulation in animal models, we have obtained preliminary clinical evidence from human specimens. Renal artery tissue samples from hypertensive patients have confirmed the altered expression pattern of miR-128 in the human renal vascular system, consistent with our findings in SHRs.

This clinical observation bridges the gap between bench and bedside. Looking ahead, we envision future clinical translational studies. We aim to conduct clinical trials investigating the therapeutic potential of siRNA-mediated silencing of miR-128 in hypertensive patients. Local administration of miR-128-specific siRNAs via renal artery intervention is expected to block the miR-128/PPAR-γ axis, thereby inhibiting the progression of hypertension and vascular remodeling. Such strategies may represent a novel targeted therapeutic approach for the management of hypertensive vascular disease[9].

Nonetheless, the findings of the present study provide a solid mechanistic framework for understanding the vascular benefits of RDN and lay a foundation for future clinical translation.

## 5 CONCLUSIONS

In summary, RDN effectively reduces renal sympathetic nerve activity and blood pressure in SHRs, and significantly alleviates hypertension-induced vascular remodeling, restores vascular diastolic function, and improves arterial baroreflex sensitivity, with a therapeutic effect comparable to that of sacubitril/valsartan

(Sac/Val). Mechanistically, RDN exerts its vascular protective effects by down-regulating the expression of miR-128, which further up-regulates the level of its target gene PPAR-γ, thereby inhibiting the overactivation of the TLR4/NF-κB inflammatory signaling pathway, reducing the release of pro-inflammatory cytokines, and suppressing vascular excessive inflammatory injury. This study clarifies that the miR-128/PPAR-γ/TLR4/NF-κB axis is a key molecular mechanism underlying the improvement of hypertensive vascular remodeling by RDN, and identifies miR-128 as a potential novel therapeutic target. These findings provide new experimental evidence and theoretical support for optimizing the clinical application of RDN and developing targeted therapeutic strategies for hypertensive vascular remodeling.

## Acknowledgments

The authors extend their gratitude to the participants enrolled in the Transcriptomic sequencing RNA sequencing analysis and to those who contributed to the publicly available data sets utilized in the present study. In addition, the authors acknowledge the investigators who facilitated data accessibility.

## Supplemental Material

## Nonstandard Abbreviations and Acronyms

Abbreviation: Full Name
miR-128: microRNA-128
PPAR-γ: Peroxisome proliferator-activated receptor gamma
TLR4: Toll-like receptor 4
NF-κB: Nuclear factor kappa B
SHR: Spontaneously hypertensive rat
RDN: Renal denervation
VSMCs: Vascular smooth muscle cells
3’UTR: 3’ untranslated region
IL-1β: Interleukin-1 beta
IL-6: Interleukin-6
TNF-α: Tumor necrosis factor alpha
TEM: Transmission electron microscopy
ICP-MS: Inductively coupled plasma mass spectrometry
IF: Immunofluorescence
IHC: Immunohistochemistry
WB: Western blotting
RNA-seq: Ribonucleic acid sequencing

